# Roots of Wheat and Rice maintain Gravitropic Setpoint Angles

**DOI:** 10.64898/2026.01.19.700323

**Authors:** Ryan Kaye, Fay Stemp-Walsh, Zoë Kitching, Alex King, Marta Del Bianco, Suruchi Roychoudhry, Stefan Kepinski

## Abstract

Root growth angle is a key determinant of root system architecture, nutrient capture efficiency and therefore yield. Yet the mechanisms governing non-vertical growth in cereal roots remain poorly understood. Here, we investigated if cereal roots maintain Gravitropic Setpoint Angles (GSAs) and the hormonal regulatory processes underpinning GSA maintenance in cereals.

Firstly, we found that both wheat seminal roots and rice crown roots actively return toward their original growth angles following displacement, consistent with true GSA maintenance. Next, we show that removal of a stable reference to gravity through clinorotation resulted in a characteristic outward curvature in all root types, indicating the presence of an antigravitropic offset similar to that described in *Arabidopsis*. Exogenous auxin treatment induced steeper rooting in both species, suggesting conserved hormonal regulatory mechanisms of GSA in both monocots and dicots. Interestingly, lateral root GSAs displayed species-specific differences: wheat laterals returned to their GSAs more effectively than rice laterals, which showed slower and incomplete responses.

Together, these findings establish that cereal roots maintain GSAs through gravity-dependent and auxin-regulated mechanisms, providing a novel framework for understanding and manipulating root system architecture in monocot crops.

## Introduction

Plant root systems are highly plastic and shaped by the integration of information from their surrounding environment. Plant organ developmental responses guided by directional stimuli are known as tropisms (Muthert et al., 2020). Plants use the constant stimulus of gravity to guide the development of both shoot and root architecture via gravitropism. Root statocyte cells contain dense starch-filled amyloplasts known as statoliths located within the gravity sensing columella cells of the root cap. In roots of flowering plants, including Arabidopsis, wheat and rice, the sedimentation of statoliths to the bottom of cells in the direction of gravity triggers the generation of an asymmetric auxin gradient, inhibition of cellular elongation and, eventually, root bending (Su et al., 2017; Roychoudhry et al., 2025).

Although the gravity vector is unidirectional, plant organs can grow at spatiotemporally determined angles, called gravitropic setpoint angles (GSAs), that can be either vertical or oblique with respect to gravity. These angles are actively maintained (Digby and Firn 1995; Hangarter and Mullen 2003; Roychoudhry et al., 2013; 2023) meaning that a root displaced from its GSA will respond by returning to its original growth angle through differential growth. Non-vertical root growth is an important adaptation to allow plants to efficiently assimilate resources: roots branch out into the surrounding soil environment to enable the plant to capture heterogeneously distributed soil resources including water, nitrogen and phosphorus over a large surface area (Lynch, 2013; Roychoudhry et al., 2017).

The monocotyledonous root systems of wheat and rice consist of embryonic roots, nodal roots, and lateral roots that can form from both root types. Wheat root system development starts with the emergence of three to six embryonic seminal roots at non-vertical angles (Richards and Passioura, 1981). Unlike the vertical primary root of Arabidopsis, these embryonic roots emerge “plagiogravitropically” or at oblique/non-vertical angles (Rich et al., 2013). The root angle of wheat seminal roots is shown to be positively associated with nodal root angle (Manschadi et al., 2008). Nodal roots develop post-embryonically from stem nodes and form most of the mature wheat root system. Lateral roots can initiate from endodermis and pericycle cells in both seminal and nodal roots (Orman-Ligeza et al., 2014). In rice, the embryonic seminal root displays a vertical growth, but is short-lived and eventually replaced by the more dominant, crown (nodal) roots (Rich et al., 2013). Nodal roots and lateral roots, which can branch off from either the seminal root or the nodal roots, develop at varied angles (Abe et al., 1994; Inukai et a, 2005). Whether these non-vertical growth angles in wheat and rice are true GSAs is still to be demonstrated.

A number of root angle mutants and root angle-associated quantitative trait loci (QTLs) have been identified in cereal species (Kirschner et al., 2024). For example, the *DEEPER ROOTING 1* (*DRO1*) and *SOIL SURFACE ROOTING 1* (*qSOR1*) QTLs both act in the regulation of root system architecture in rice (Uga et al., 2013; Kitomi et al., 2020). *DRO1* homeologs have been identified and found to have root tip expression in wheat (Ashraf et al., 2019). *DRO1* and *qSOR1* belong to the *LAZY* family, which play a crucial role in plant gravitropism and the regulation of both shoot and root architecture (Jiao et al., 2021). *ENHANCED GRAVITROPISM1 (EGT1) and ENHANCED GRAVITROPISM2* (*EGT2*) can regulate root growth angle in barley and wheat lines. Plants carrying mutations for either genes show steeper seminal and lateral root angle (Fusi et al., 2022, Kirschner et al., 2021). While this evidence collectively suggests that non-vertical growth angles are genetically set in monocots, true root GSAs must be maintained in respect to the gravity vector.

Over the last decade, studies have demonstrated that, as well as regulating gravitropism, auxin plays a crucial role in the GSA maintenance. In the model plant *Arabidopsis thaliana*, it has been proposed that two opposite auxin fluxes, determine the gravitropic response and an antagonistic antigravitropic offset (AGO) (Roychoudhry et al., 2013). The counteraction of the AGO to the gravitropic response allows for the setting and maintenance of the GSA. In Arabidopsis, the AGO is proposed not to be active in primary roots, resulting in vertical growth (Roychoudhry et al., 2013; 2023). Primary roots can skew from the vertical due to mechanical interaction between the root tip, rotating about its axis, and the substrate (Vaughn and Masson, 2011). To be considered true GSAs, non-vertical root angles should however be maintained through an AGO or an equivalent mechanism.

Here, we investigate if monocot root systems actively maintain GSAs. We also use simulated microgravity via 2D clinorotation to determine if non-vertical roots possess an AGO and further test the effect of auxin on root GSA in monocots. Our findings provide the first molecular framework to the understanding of mechanisms that regulate non-vertical growth patterns in roots of monocot species.

## Results

### Wheat seminal roots and rice crown roots actively maintain gravitropic setpoint angles

The first step to establish whether non-vertical growth angles of cereal roots are true GSAs is testing if they are maintained after a change in orientation with respect to gravity. To this aim, we performed reorientation experiments with wheat cv. Bobwhite seminal roots and rice cv. Nipponbare crown roots. Plants were grown vertically and then reorientated by 30° for 24 hours with the root tip angles measured before and after reorientation. A 30° reorientation angle was chosen so roots would not be reorientated past the vertical axis (Figure 1). A 30° reorientation angle also meant that roots were either reorientated above (downward bending) or below (upward bending) their growth angles. Wheat seminal roots returned to their original angle when downwards bending (Figure 1a, white arrows, and c). Upwards bending wheat seminal roots bent toward the angle (Figure 1a, black arrows), but were still more vertical than the pre-reorientation angle after the 24 h period (Figure 1c). Similarly, rice crown root also showed faster downwards bending than upward bending (Figures 1b, d). This could suggest that wheat seminal roots and rice crown roots actively maintain their growth angles, implying that their non-vertical growth angles could fall into the definition of GSAs.

**Figure 1:**
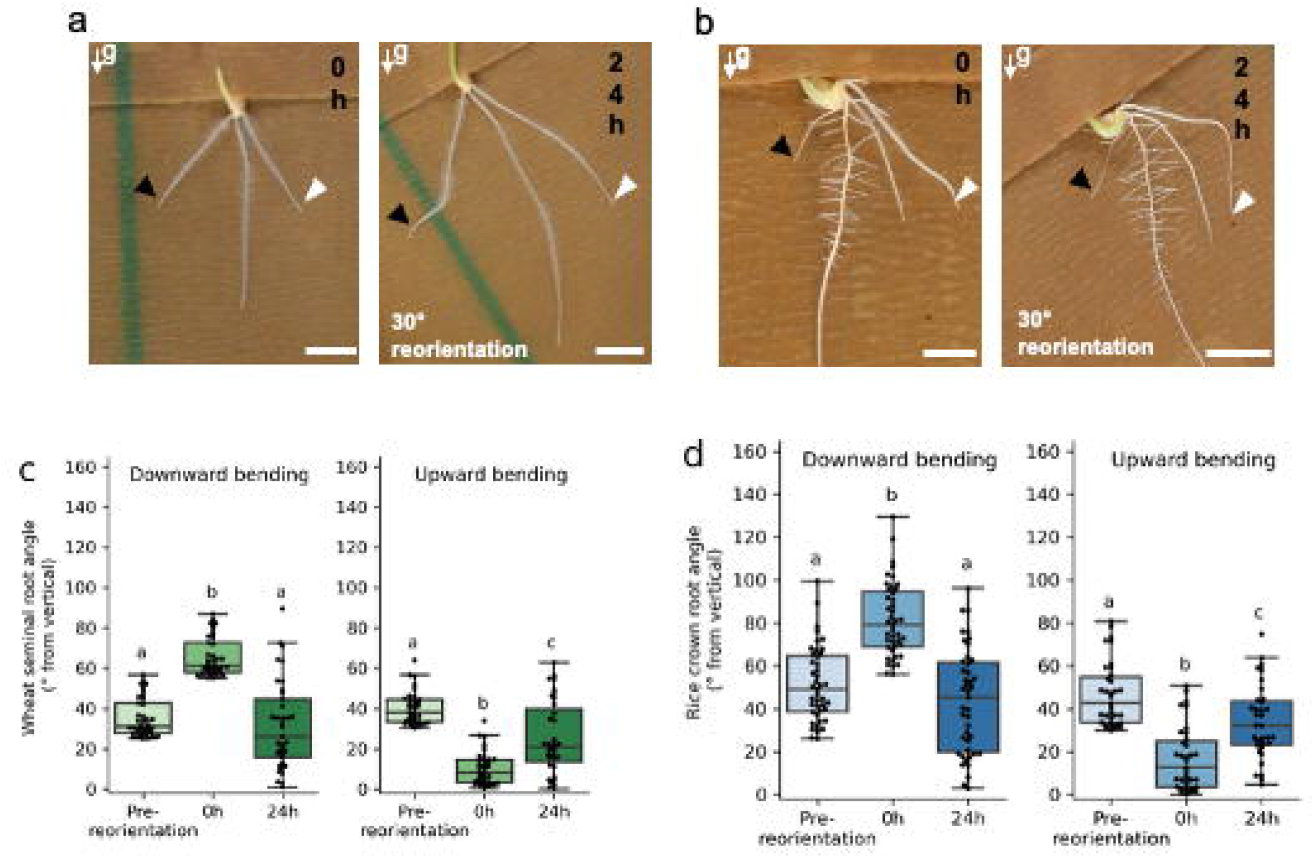
Wheat seminal and rice crown roots return towards their original angle after reorientation. Upward (black arrows) and downward (white arrows) bending wheat ‘Bobwhite’ seminal (a, c) and rice ‘Nipponbare’ crown (b, d) roots reorientated at 30° for 24 hours. Plants growth in germination pouches and and 24 hours, ‘g’ = direction of gravity, scale bar = 10 mm, p<0.05.

### Wheat seminal and rice crown roots bend outwards under clinorotation

Because the AGO, which balances the gravitropic response, is stable in the timeframe of the graviresponse, the removal of a stable reference to gravity, such as by 2D clinorotation, induces a characteristic outward curvature in Arabidopsis lateral roots (Roychoudhry et al., 2013). We therefore used clinorotation to investigate whether wheat seminal and rice crown roots would display the sign of an AGO in response to removal of gravity input. After 7 hours of 1 rpm clinorotation, outward and upward curvature of wheat seminal and rice crown roots was seen, similarly to what was observed in Arabidopsis lateral roots (Figure 2, black arrows). Interestingly, in contrast to vertical Arabidopsis primary roots, which do not display any changes after 2D clinorotation (Roychoudhry et al., 2013), vertically growing wheat and rice seminal roots displayed upward bending similar to crown roots (Figure 2, white arrows). We hypothesize cereal seminal roots may possess an anti-gravitropic offset that enables growth at a slight non-vertical growth angle (Figure 2e). In this context, it is likely that the three wheat seminal roots arrange themselves as pyramid, that gets flattened on the 2D experimental surface (Supplementary Figure 1).

**Figure 2:**
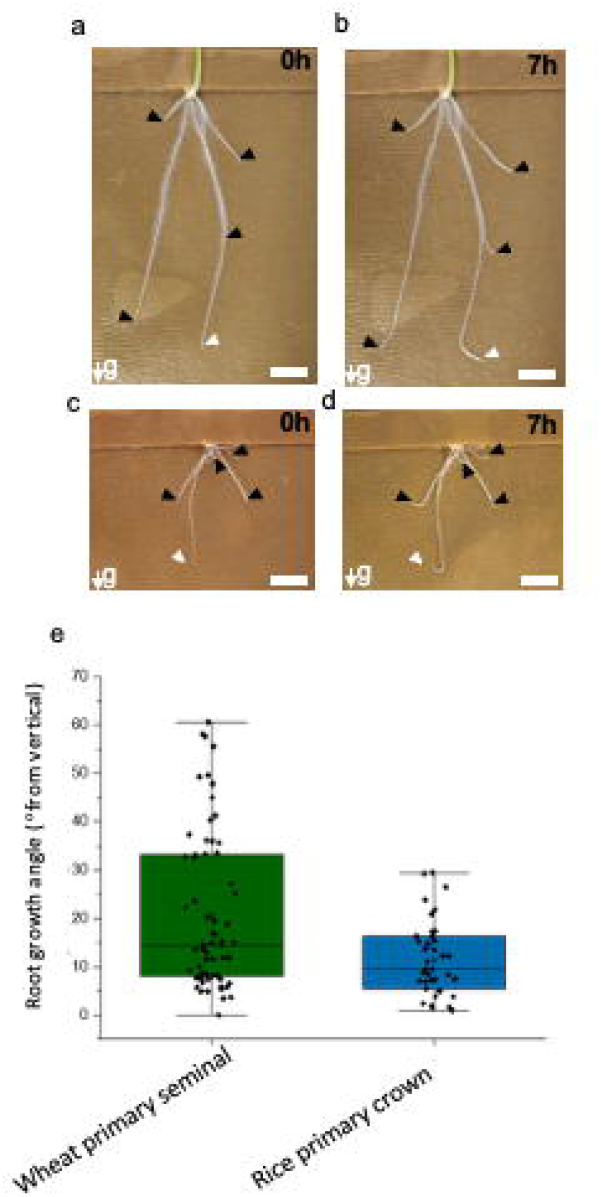
Wheat seminal roots and rice crown roots bend outward upon clinorotation. Wheat seminal roots (a-b) and rice crown roots (c-d) pre-clinorotation (a, c) and root responses to clinorotation after 7 hours (b-d) Arrows indicate root tips, with white arrows showing primary root tips. Scale bars= 10 mm, ‘g’ represents the direction of gravity pre-clinorotalion. (e) Quantification of primary seminal and crown root growth angles in wheat and rice seedlings.

### Auxin induces deeper rooting in wheat and rice

Non-vertical GSA is regulated, in Arabidopsis seedlings, by the plant hormone auxin (Roychoudhry et al., 2013; Ruiz Rosquete et al., 2013). We therefore tested if auxin was able to influence the root growth angle of wheat seminal and rice nodal roots. We treated wheat and rice seedlings with increasing concentrations of the naturally occurring auxin indole-3-acetic acid (IAA). IAA treatment made both, wheat seminal and rice crown roots more vertical (Figure 3). However, root growth angle of both species did not change significantly across the range of auxin concentrations tested, indicating that the effect of auxin on cereal root growth angle is not dose-dependent in the investigated hormone range. By showing that auxin also regulates root growth angle in wheat and rice, our findings could suggest that similar molecular mechanisms regulate GSA in Arabidopsis and cereals.

**Figure 3:**
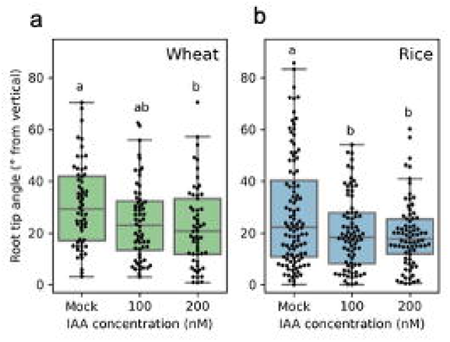
Wheat seminal and rice crown root tip angles with IAA treatment. Wheat (a) and rice (b) root tip angles after treatment with 100 nM and 200 nM IAA, p<0.05.

### Wheat and rice lateral roots demonstrate differences in GSA maintenance

In cereals, both seminal and grown roots can produce non-vertical lateral roots (Kirschner et al., 2024). Initial analysis of lateral root GSAs of wheat and rice showed species-specific differences, with wheat having a steeper GSA than the more horizontally emerging rice lateral roots (Supplementary Figure 2). To assess whether lateral root growth angles in wheat and rice were also maintained relative to gravity, wheat and lateral roots were subjected to the same tests used on wheat seminal and rice crown roots,

First, wheat and rice lateral roots were gravistimulated at 30° (Figure 4 a-d). In wheat, similarly to seminal roots, lateral roots showed a faster downward bending and a slower upward bending, and were able to return to their original growth angles or close, respectively. In rice lateral roots, instead, both the upward and downward bending response were very slow, and roots failed to return to their original growth angles after 24 hours. To assess if longer time would allow cereal lateral roots to return closer to their original GSA, we performed reorientation experiments over 48 hours (Supplementary Figure 3). We found that, in rice, downwards bending lateral roots never returned to their original GSA and only displayed a small change in angle. Upwards bending rice laterals roots, instead, returned closer towards their GSA after 36 hours with a wide range in angles (Supplementary Figure 3b). Interestingly, while both upward and downward bending wheat lateral roots returned to their GSA, after only 12 hours from reorientation they became progressively more vertical at successive timepoints (Supplementary Figure 3a).

**Figure 4:**
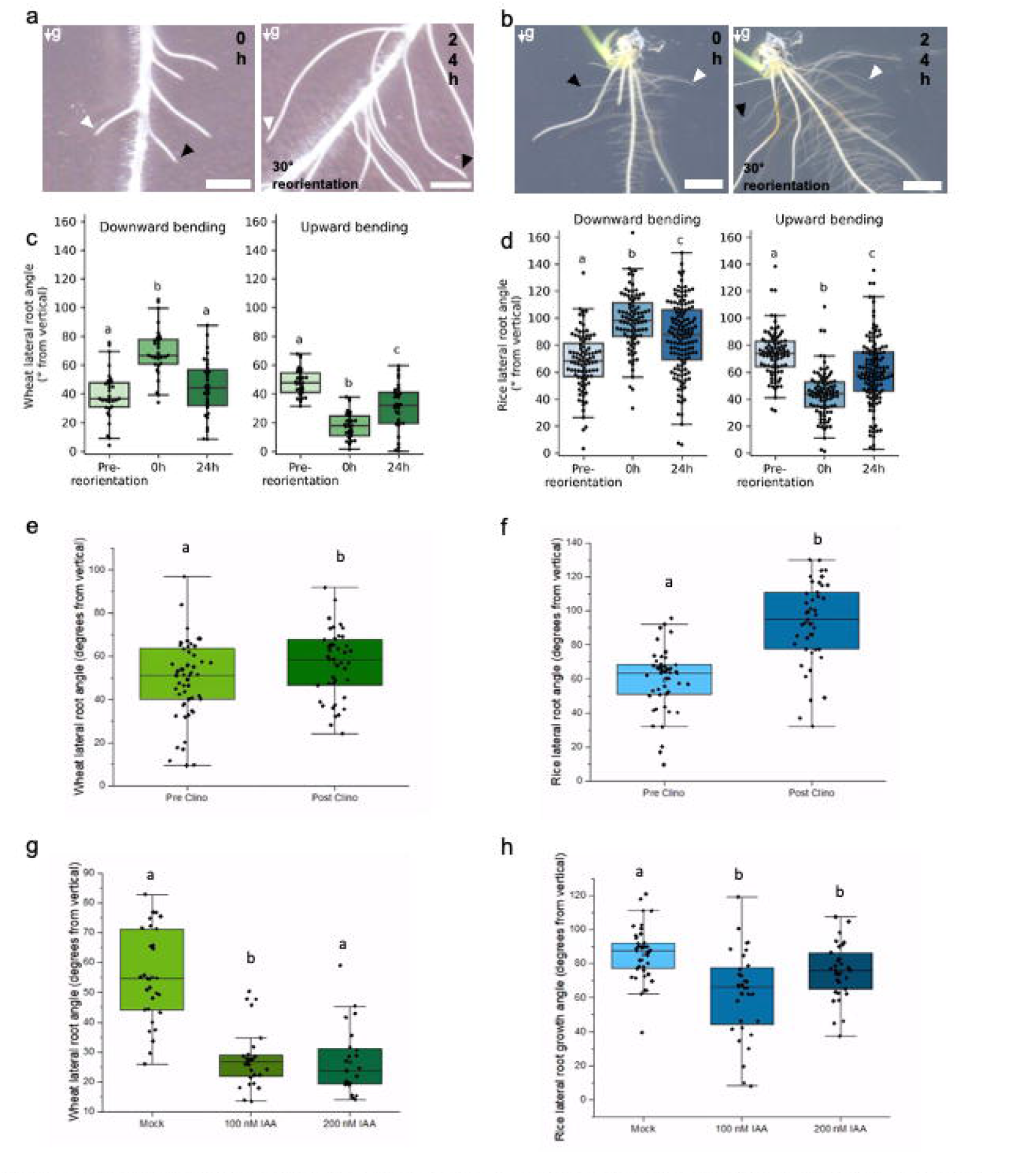
Wheat and rice lateral root growth angle response after 30° reorientation. Upward (black downward (white arrows) bending wheat ‘Bobwhite’ (c) and rice ‘Nipponbare’ (d) lateral roots reoriented at 30° for 24 hours. Plants grown on agar and imaged at 0 and 24 hours, ‘g’ = direction of gravity, mm, p<0.05. Change in angle in wheat (e) and rice (f) latera roots post 6 hours of clinorotation at 4 r. p. m. Both, wheat and rice lateral roots show significant and characteristic upward and outward bending lion. Effect of auxin treatment on lateral root GSA in wheat (g) and rice (h) lateral roots. Auxin induces steeper rooting in lateral roots of both species.

Next, wheat and rice lateral roots were tested for the presence of an AGO and for their responsiveness to auxin in the context of the control of non-vertical growth. Clinorotation was used to remove a stable reference to gravity to investigate whether wheat and rice lateral roots would display the sign of an AGO. After 6 hours of clinorotation, both wheat and rice lateral roots displayed the characteristic upward bending (Figure 4e, f). While wheat lateral roots bent upwards to a similar degree as wheat seminal roots, the upward bending in rice lateral roots was very prominent compared to rice crown roots. Conversely, while the growth angle regulation by auxin in rice laterals (Figure 4h) was comparable to that of crown roots, growth angles of wheat lateral root were much more responsive to auxin treatment than any other root type investigated (Figure 4g). Interestingly, while auxin induced steeper rooting in rice lateral roots at both, 100 and 200 nM, wheat lateral roots became shallower at 200 nM of auxin treatment. These data highlight not only the difference in growth angle maintenance between cereal species, but also between root types.

### Seminal and lateral roots of wheat and rice gradually adopt more vertical orientations in developmental time

To assess whether any of the observed phenotypes in response to gravistimulation could be dependent on a change in growth angle in time, as observed in Arabidopsis lateral roots (Mullen and Hangarter, 2003), we characterised root growth angle over time in different root types of wheat and rice. Wheat seminal roots and rice crown roots form at similar stages in plant development and emerge at non-vertical angles, in a similar manner to the lateral roots of these species. The root growth angle from the vertical axis was measured for wheat seminal and rice crown roots in 5-day old plants. The seminal growth angle became gradually more vertical during development with the angle of rice crown roots being steeper than wheat seminal roots (Figure 5). Since lateral roots emerge 6-10 days post germination in monocots (Wang et al., 2018), plants were grown for 10 days prior to analysis of lateral root development. Lateral roots of wheat became slightly more vertical over time compared to rice lateral roots, which maintained a more horizontal angle throughout the time period.

**Figure 5:**
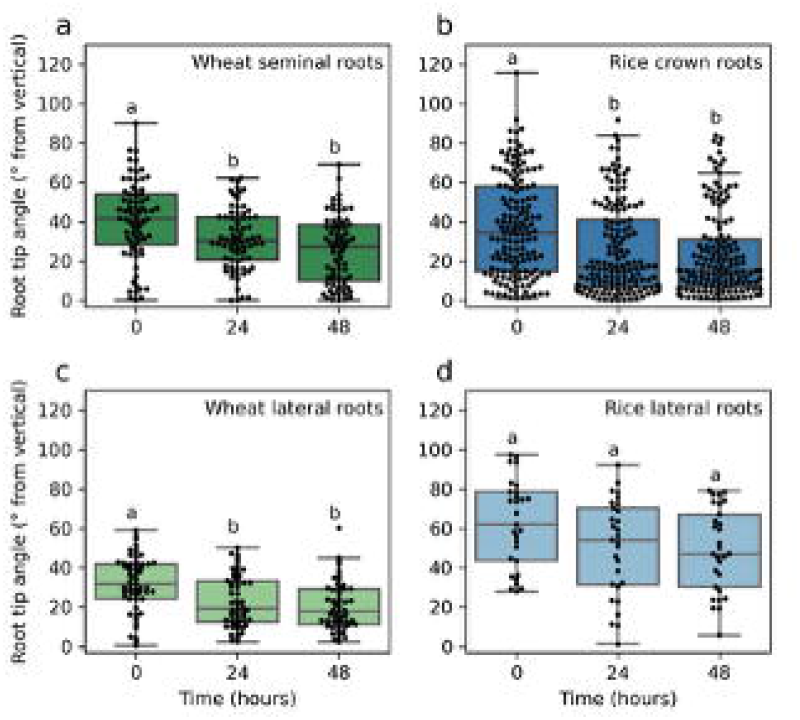
Change in wheat and rice root growth angles over time. Wheat seminal (a), rice crown (b), wheat lateral (c) and rice lateral (d) root tip angles over 48 hours, p<0.05.

## Discussion

The establishment of non-vertical growth angles is the main determinant of plant architecture. Understanding the mechanisms of root growth angle control in crop species is therefore important for optimising root system architecture. Root GSAs have been well characterised in Arabidopsis lateral roots but less so in other monocot species (Roychoudhry et al., 2013; 2023, Ruiz-Rosquete et al.,2013, 2017). Here, we found that the non-vertically growing roots of wheat and rice maintain gravitropic setpoint angles that change during root development and in response to loss of a constant gravity stimulus or application of exogenous auxin. Our findings give insight into the mechanisms for cereal root growth angle control and show differences between wheat and rice monocot root systems.

The fibrous root system architecture of cereals differs dramatically from that of the dicot Arabidopsis, so we used the early-forming and non-vertical wheat and rice roots for GSA analysis. Our results showed that all wheat and rice root types investigated became more vertical over a 48-hour period with wheat seminal and rice crown roots becoming steeper in the first 24 hours, whereas lateral roots showed an altered GSA between 24 and 48 hours. Wheat lateral roots had a faster change in GSA over time than in rice. Wheat lateral roots have an initial steeper GSA than rice lateral roots which were less vertical. Wheat seminal roots have been described to change to a more downwards orientation with distance from the base of the plant which could be when roots reach an older developmental stage. The authors hypothesised this could be due to internal or in combination with external factors controlling the root system development (Nakamoto et al., 1994). This shows the importance of the genetic and environmental regulation of GSA maintenance and root gravitropic responses. Rice crown roots also show plagiotropic growth and it has been proposed that the crown root growth angle is determined by its emergence position as well as a potential correlation between root diameter and the gravitropic response (Oyanagi et al., 1993).

Root responses to the change in gravity stimulus from a 30° reorientation differed between the two species. Wheat seminal and rice crown roots both returned towards their GSAs for both upwards and downwards bending roots. Wheat lateral roots returned to their original GSA faster and then became more vertical in comparison to rice laterals which were still returning towards their GSAs after 48 hours. This could be an indicator of a difference in root growth rate or the scale of gravitropic response between wheat and rice root systems. Rice only forms one seminal root so more horizontal lateral roots may allow coverage of a larger surface area. Comparisons to wheat lateral root GSA maintenance can be made with Arabidopsis lateral roots as both have similar lateral root GSAs which are steeper than rice (Roychoudhry et al., 2013), so rice could potentially have more differences in root gravitropic responses. The differences in GSAs and reorientation root growth responses indicate that gravitropic response mechanisms differ within different monocot species as well as in comparison to dicots. Arabidopsis mutants with higher levels or auxin or increased auxin response have more vertical lateral roots than wildtype (Roychoudhry et al., 2013), so it is possible rice lateral roots could potentially have lower auxin levels or a stronger auxin-dependent AGO than in wheat lateral roots.

Clinorotation results in omnilateral gravitational stimulation and this loss of a constant gravity reference causes an outwards and upwards bending growth response in Arabidopsis lateral roots (Roychoudhry et al., 2013). Observations following wheat and rice clinorotation found similar root responses in wheat seminal and rice crown roots. The first wheat seminal root to form (“the primary seminal root”) usually grows vertically in a 2D growth system as seen here, however, this root was also observed to grow up- and outwards with clinorotation. A study of wheat cultivars grown in the soil in field conditions found the primary seminal root does not grow vertically but at a smaller angle than the other later-forming seminal roots (Nakamoto et al., 1994), so all wheat seminal roots are likely to have an AGO unlike the vertically growing Arabidopsis primary root.

A study of plagiotropic or non-vertical wheat seminal roots are suggested to be influenced by the external environment and by internal factors including growth hormones (Nakamoto et al., 1994). Arabidopsis lateral root GSA is auxin-regulated and auxin treatment causes more vertical lateral root growth (Ruiz Rosquete et al., 2013; Roychoudhry et al., 2013) . Arabidopsis lateral roots shift to more vertical GSAs with 50-100nM IAA treatment and a similar finding is also seen with bean basal and lateral roots (Roychoudhry et al., 2017). We found that IAA treatment increased the verticality of both wheat seminal and rice crown root GSA, indicating auxin is important in GSA maintenance for these cereal species. Auxin may be a factor in the differences seen between the lateral roots of wheat and rice. One initial study of the effects of IAA treatment on wheat seminal roots showed only small effects on root growth angle but a decrease in root elongation growth rate for two different wheat cultivars (Oyanagi et al., 1993). More recent findings in barley cv. ‘Morex’ showed barley had significantly steeper seminal root angles following IAA treatment and 90º reorientation (Fusi et al., 2022). Auxin (IAA) application shown to restore root gravitropic response in rice lateral rootless mutants (*Lrt1*) of rice showing auxin is important in rice root gravitropism (Chhun et al., 2003). Studies have shown that wheat and rice have many gene families important for auxin function including *auxin response factor* (*ARF*) genes (Qiao et al., 2018; Sato et al., 2001) and *PIN-FORMED* (*PIN*) genes (Kumar et al., 2021).

Knowledge of root angle control in crop species provides a basis for altering root phenotypes to improve nutrient uptake as different root system ideotypes are proposed to optimise resource capture for water and different nutrients. The ‘steep, cheap and deep’ ideotype of more vertical root systems can increase nitrogen and water uptake, whereas a shallower topsoil rooting ideotype could improve phosphorus capture (Lynch, 2019). Indeed, maize roots become steeper when grown nitrogen deficient field conditions (Trachsel et al., 2013). Moreover, a study of Japanese wheat cultivar seminal root angles under controlled conditions showed northern cultivars were deeper-rooting than the southern cultivars which are usually grown under wetter conditions (Oyanagi et al., 2003). *DRO1* was first identified in rice and functions in root growth angle control as *DRO1* overexpression leads to more vertical root systems and higher yields in water-limited environments (Uga et al.,2013). In rice, the loss of function of a *DRO1* homolog named *qSOR1* (*quantitative trait locus for SOIL SURFACE ROOTING 1*) resulted in shallower rooting rice which improved yield under saline conditions (Kitomi et al., 2020). Characterisation of root system ideotypes and discoveries of root angle regulating genes in crop species gives potential for optimising root system architecture (Uga, 2021). Understanding of root gravitropic responses in cereal species is important as the ability to alter root systems for environmental adaptation is key for future crop yield increases.

## Materials & Methods

### Plant Materials

Wheat ‘Bobwhite’ and rice ‘Nipponbare’ varieties were used for all experiments. Wheat experiments were grown under conditions of 22ºC day, 15ºC night, 16-hour photoperiod and rice experiments were grown in 27ºC, with a 12-hour photoperiod. Seeds were surface sterilised for 3 hours using chlorine gas seed sterilisation. Wheat ‘Bobwhite’ seeds or de-husked ‘Nipponbare’ rice seeds were placed on moist filter paper and cold treated at 4ºC for 2 days before use. Rice seeds were then additionally moved to 27ºC, 12-hour photoperiod for 2 days to allow germination.

### Wheat seminal and rice crown root reorientations

Seeds were placed into Cyg seed germination pouches (Mega International, Minneapolis, US) with the embryo orientated so that the germ was facing outwards and downwards. Pouches were wrapped in foil to exclude light and placed upright in a reservoir of Hoagland’s No2 basal salt solution (Hoagland and Arnon, 1950) and plants were allowed to grow for 5 days (wheat) or 2 days (rice). Plants were imaged, reoriented by 30º, and reimaged after 24 hours. Seminal root tip angles were analysed before and after reorientation using RootNav (Pound et al., 2013) and measurements processed using Microsoft Excel and Python. Roots that showed no growth or with a starting angle below 30º were excluded from the analysis as reorientation would have placed the roots beyond the vertical (0°) and transposed them to the other side. All images were captured using a Sony Cyber-Shot DSC RX100. Non-reorientated control plants were imaged at the same time as the reorientated plants but not reorientated.

### Wheat and rice lateral root reorientations

10-day wheat and rice seedlings were grown on 245 mm (wheat) or 120 mm (rice) square plates in Hoagland’s No2 (wheat) or Yoshida’s (rice) medium (Yoshida et al., 1971) in their respective growth conditions. Plates were reorientated by 30º and imaged using an Epson Perfection V800 flatbed scanner. Lateral root tip angle measured as the angle from vertical of a 1 mm segment measured from the root tip were analysed with ImageJ for all lateral root experiments. Non-reorientated control plates were imaged at the same time as the reorientated plants but not reorientated.

### Wheat and rice root clinorotation

Plants were grown as above but instead of reorientation plants were placed upon a 1 rpm clinostat in their respective growth conditions. Wheat plants were clinorotated at room temperature and rice plants were clinorotated at 27°C. Plants had light excluded from the roots and were clinorotated for 7 hours with imaging before and after clinorotation.

### Wheat and rice auxin treatments

Plants were grown as for reorientations but were placed in a reservoir of Hoagland’s No 2 containing IAA (or mock treated) to a given concentration from a 100 mM stock solution of IAA dissolved in 70% ethanol. Wheat plants were photographed after 6 days of growth and rice plants were photographed after 4 days of growth in the pouches. For lateral root auxin treatments, plants were grown as described above, but for 10-12 days to allow lateral roots to develop. Plants were not reorientated and tip angles of wheat seminal roots and rice crown roots were measured with RootNav (Pound et al., 2013).

## Author contributions

R.K and F. S-W. did most of the experiments and wrote the manuscript except: Z.K. provided the rice and wheat clinorotation data. A.K. imaged wheat lateral root systems. S.R. generated the seminal and primary root angle data. S.K conceptualised the study and provided final edits to the manuscript with S.R and M.D.B.

